# Guggulsterone Enhances NKX3.1 Expression and Induces Apoptosis in Prostate Cancer Cells: Implications for Chemoprevention

**DOI:** 10.1101/2023.12.08.570071

**Authors:** Garima Jain, Neha, Prashant Ranjan, Chandra Devi, Aditi Atri, Clara Cieza-Borrella, Parimal Das

## Abstract

**Aim:** Prostate cancer is a leading cause of cancer-related mortality, necessitating novel therapeutic and preventive strategies. This study aimed to investigate the anti-cancer properties of Guggulsterone (GS) on prostate cancer cells, specifically focusing on its effects on homeobox protein NKX3.1, apoptosis, and mitochondrial function.

**Method:** The study utilized the LNCaP prostate cancer cell line and involved cell culture, dose-response analysis, flow cytometry for apoptosis quantification, siRNA-mediated knockdown of NKX3.1, mitochondrial membrane potential assessment using TMRM staining, qRT-PCR analysis, and Western blotting. The stability of NKX3.1 mRNA and protein in response to GS was also evaluated.

**Result:** GS treatment induced apoptosis in LNCaP cells in a dose and time-dependent manner, with an impact on early and late-stage apoptosis. This effect was mediated, at least in part, through the upregulation of NKX3.1. GS enhanced NKX3.1 expression at both the protein and mRNA levels, even in the absence of androgens. Furthermore, GS treatment extended the half-life of NKX3.1 mRNA and protein, suggesting an effect on their stability.

**Conclusion:** This study provides valuable insights into the anti-cancer mechanisms of GS in prostate cancer cells. The upregulation of NKX3.1 by GS and its role in mediating apoptosis offers NKX3.1 as a potential prevention target for PCa. The findings open new research avenues for the development of targeted therapies that stabilize NKX3.1 expression and protect mitochondrial function. Further investigations are needed to understand the intricate molecular mechanisms underlying GS’s effects fully, potentially improving prostate cancer management and outcomes.

## Introduction

Prostate cancer (PCa) is a potent malignancy predominantly affecting older males and remains a leading factor in global cancer-related mortality (1). Its impact goes beyond fatality rates to include substantial economic and social costs related to diagnosis, treatment, and long-term care (2). Traditionally, the primary focus in combating PCa has revolved around early detection and a range of treatment modalities, from chemotherapy to hormonal and radiation therapy that suffer from therapeutic resistance or present considerable side effects (3) necessitating the need for preventive therapies. More recently, the molecular and genetic aspects of the disease, especially the role of the homeobox protein NKX3.1, have come under scrutiny (4). Research has established NKX3.1 has a vital role in PCa initiation and progression (5,6), as well as a possible marker for prostatic origin. NKX3.1 plays role during normal prostate gland development, where it is expressed primarily in luminal epithelial cells, which are responsible for the production and secretion of prostatic fluid. Its expression is tightly regulated and exhibits a highly restricted pattern, with minimal expression observed in other tissues. This spatial and temporal specificity of NKX3.1 expression highlights its importance in maintaining the normal function and integrity of the prostate gland. Additionally, NKX3.1 controls the expression of genes involved in cellular differentiation, secretory function, and androgen receptor signaling, which are critical for the normal function of the prostate gland (4,7).

The reduced NKX3.1 expression is considered an early event in prostate tumorigenesis, occurring even before the development of histological abnormalities in the prostate gland (8). The loss of NKX3.1 expression in PCa can be attributed to various mechanisms, including genetic alterations, epigenetic modifications, accelerated protein degradation, and dysregulation of signaling pathways. Loss of NKX3.1 in PCa disrupts the balance of these signaling pathways, contributing to the acquisition of a more aggressive and dedifferentiated phenotype, leading to uncontrolled cell proliferation (9). This results in prostatic epithelial dysplasia and benign hyperplasia. Understanding the precise molecular mechanisms underlying NKX3.1 dysfunction in PCa may provide insights into novel therapeutic and preventive strategies; for instance, rescuing NKX3.1 expression could serve as an effective strategy for chemoprevention. Nonetheless, the field lacks targeted interventions capable of harnessing molecular understanding of NKX3.1 to halt disease progression effectively.

In recent years, natural compounds and traditional medicines have gained attention for their potential in cancer prevention and treatment (10). One such compound is Guggulsterone (GS), a non-toxic bioactive compound derived from the resin of the guggul tree (*Commiphora wightii*), which has been used for centuries in traditional Ayurvedic medicine. GS exists in two isoforms naturally - E and Z. z-GS possesses various pharmacological properties, including anti-inflammatory and antioxidant activities (11). Z-GS has demonstrated significant pharmacological effects that demonstrate its efficacy as a strong anticancer drug (12). In PCa cells, GS has been shown to induce apoptosis (13). It is known that GS affects the action of several nuclear receptors, growth factors, and transcription factors at the molecular level (14,15). But there is still much to learn about the exact sequence of GS’s underlying anticancer process. The most well-established mechanism for GS’s anti-cancer activity has been linked to the endpoints of the apoptotic cascade, which includes pro-apoptotic pathways (JNK) being activated (16,17), anti-apoptotic factor (NF-KB) being inhibited (18), and pro-apoptotic factors BAX-BAK being increased (19). While GS’s impact on important molecular targets (like NF-kB, PI3K/AKT, JAK/STAT, c-Myc, MAPK, VEGFR, and Wnt/b-catenin) in a variety of cancers (like head and neck, colon, breast, prostate, pancreatic, and hepatocellular carcinoma) has already been evaluated (20–24), no study has examined GS’s impact on pathways and proteins specific to PCa pathogenesis, like NKX3.1 loss of expression, ERG-TMPRSS2 fusion, expression and stability of androgen receptor (AR) cofactors, EZH2 over-expression, AR synthesis enzymes, growth factor receptor activity, and activity of hormone-chaperons like androgens-HSP90 (25).

Our study is the first to investigate z-GS’s effects (z-guggulsterone referred to as GS hereafter) on unique molecular pathways and genetic markers specific to PCa, like NKX3.1, in a time- and dose-dependent manner. These steps introduced a novel layer of understanding to the involvement of NKX3.1 in GS-induced apoptosis, in the context of PCa. We used PCa cell lines in this study and we have demonstrated that GS-induced apoptosis is mediated by NKX3.1. Furthermore, GS treatment effectively elevated NKX3.1 protein and mRNA expression levels in the LNCaP PCa cell line in a time- and dose-dependent manner. Additionally, the half-life of NKX3.1 mRNA and protein was prolonged in response to GS treatment. Mitochondrial membrane potential was also improved following GS treatment, further implementing that enhanced mitochondrial function, might help prevent or slow down the development PCa. However, it should be noted that GS fails to counteract inflammation-induced NKX3.1 loss, emphasizing the importance of considering the effects of traditional drugs in a tumor- and stage-specific manner. Collectively, our findings suggest that GS exhibits chemopreventive potential by enhancing NKX3.1 expression and inducing apoptosis in PCa cells

## 2. Materials and Methods

### 2.1 Cell culture, dose-response analysis, and cell treatment

PCa cell line LNCaP was cultured with Rosewell Park Memorial Institute (RPMI) medium supplemented with 10% FBS and 1X penicillin-streptomycin cocktail at 37°C in a humidified environment with 5% CO2. For each experiment, cells were seeded in a T25 flask and grown for 1 day in a complete RPMI medium, reaching 60-70% confluence. Different GS concentrations were applied to 60-70% confluent cells for the required time period. Additionally, cells were pretreated with 40 μM of GS for the experiments where cells were exposed to Actinomycin-D and Cyclohexamide.

### 2.2 Flow cytometry analysis

By double labeling with AnnexinV-APC and propidium iodide (PI), apoptotic cells were measured. Following treatment, cells were harvested by pelleting down at 3000 rpm for 3 minutes, and they were then twice washed with PBS. Following a 15-minute dark incubation period at 37°C, the cells were reconstituted in 1X binding buffer (supplied with the Annexin V-APC/PI kit), stained with 5 μl of Annexin V-APC, and then exposed to 1 μl of PI. Flow cytometry was used for the measurements (CytoFLEX LX Beckman Coulter). There were 10,000 events in each sample.

### 2.3 NKX3.1 gene knockdown

NKX3.1 expression was inhibited using siRNA against the NKX3.1 gene (WD10664038) as well as universal negative control SiRNA#1 (SIC001) was purchased from Sigma Aldrich. 70-90% of confluent cells were replaced with Opti-MEM. According to the manufacturer’s instructions, cells were transfected with 400 nM of siRNA and equimolar negative siRNA as a control using the Lipofectamine 3000 reagent. Cells were incubated for 24 hours followed by treatment with 40 μM of GS for an additional 24 hours.

### 2.4 Measurement of mitochondrial membrane potential

Tetramethylrhodamine methyl esters (TMRM) staining was used to assess mitochondrial membrane potential in response to GS treatment. After 48 hours of GS incubation, the cells were collected, washed with PBS, and then stained with 200 nM TMRM in PBS at 37°C for 15 minutes in the dark. The intensity of TMRM’s orange fluorescence was measured using flow cytometry.

### 2.5 qRT-PCR analysis

Quantitative reverse transcription-polymerase chain reaction (qRT-PCR) was used to analyze gene expression. TRIzol reagent was used to extract total cellular RNA, and DNaseI treatment was then applied to eliminate DNA contamination. Spectrophotometry (NanoDrop, Thermo, Waltham, MA) was used to measure the quantity and quality of RNA. A cDNA synthesis kit (Applied Biosystems, Waltham, MA) was used, and 1 µg of pure RNA was converted to cDNA by following the manufacturer’s instructions. Using qRT-PCR (QuantStudio 6, ThemoFisher Scientific) and the SYBR Green Real-time PCR master mix (Thermo Scientific), the expression levels of certain genes were determined.

### 2.6 Western blotting

TNT buffer was used to prepare whole-cell extracts. Protease inhibitors (COMPLETE Protease Inhibitor; Roche Applied Sciences) were added to the extraction buffers. buffers A and C were used to isolate nuclear and cytoplasmic proteins. Standard methods were followed to separate 50–100 μg of protein extract on SDS-PAGE and transfer it to nitrocellulose membranes (Schleicher & Schuell, Whatman) for immunoblot analysis. Before being incubated with the primary antibody (1:1000 in TBS-Tween-20), the membrane was blocked with 5% milk powder in TBS-Tween-20. It was then washed and incubated in a TBS-Tween-20 solution containing a secondary antibody (1:5000) coupled with horseradish peroxidase. Thermo Scientific Pierce ECL substrates were used for the detection. Using the ImageJ program, the scanned pictures were examined for densitometric quantification. TNT buffer: 20 mM Tris pH 8.0, 200 mM NaCl, 1% Triton X-100, 1 mM DTT, protease inhibitor. Buffer A: 10 mM HEPES pH 7.9, 10 mM KCl, 0.1 mM EDTA, 0.1 mM EGTA, 1 mM DTT, 0.5 mM PMSF. Buffer C: 20 mM HEPES pH 7.9, 0.4 M KCl, 1 mM EDTA, 1 mM EGTA, 1 mM DTT, protease inhibitors. anti-NKX3.1 (sc-393190), ERK2 (sc15022), anti-AR (cell signaling #3202), Rela (cell signaling #8242) and Cleaved caspase 3 (cell signaling #9664T)

### 2.7 Statistical Method

Every experiment was conducted three times, and the results were given as the mean standard deviation (mean). The paired student t-test was used for the statistical analysis. The differences were considered statistically significant when the P-value was ≤ 0.05. By plotting the time period against the NKX expression values, a linear regression curve was used to derive the T half-life.

## 3. Results

### 3.1 Guggulsterone Elicits Apoptotic Responses in Prostate Cancer Cells

To extend the current body of knowledge concerning the pro-apoptotic effects of guggulsterone (GS) in prostate cancer (PCa) cells, we rigorously examined the optimal GS concentrations and treatment durations for this investigation.

To quantify cellular viability and proliferation in response to GS administration, an MTT assay was performed on both PC3 and LNCaP cell lines across a GS concentration gradient (1, 5, 10, 20, 40, and 60 µM) (Fig 1A). The calculated IC50 for LNCaP cells stood at 43 µM, signifying a notable reduction in cellular viability at this concentration (Figure 1A). In contrast, PC3 cells exhibited no statistically significant variations in cell viability across the examined concentration spectrum (Supplementary Figure S1a).

**Figure 1:**
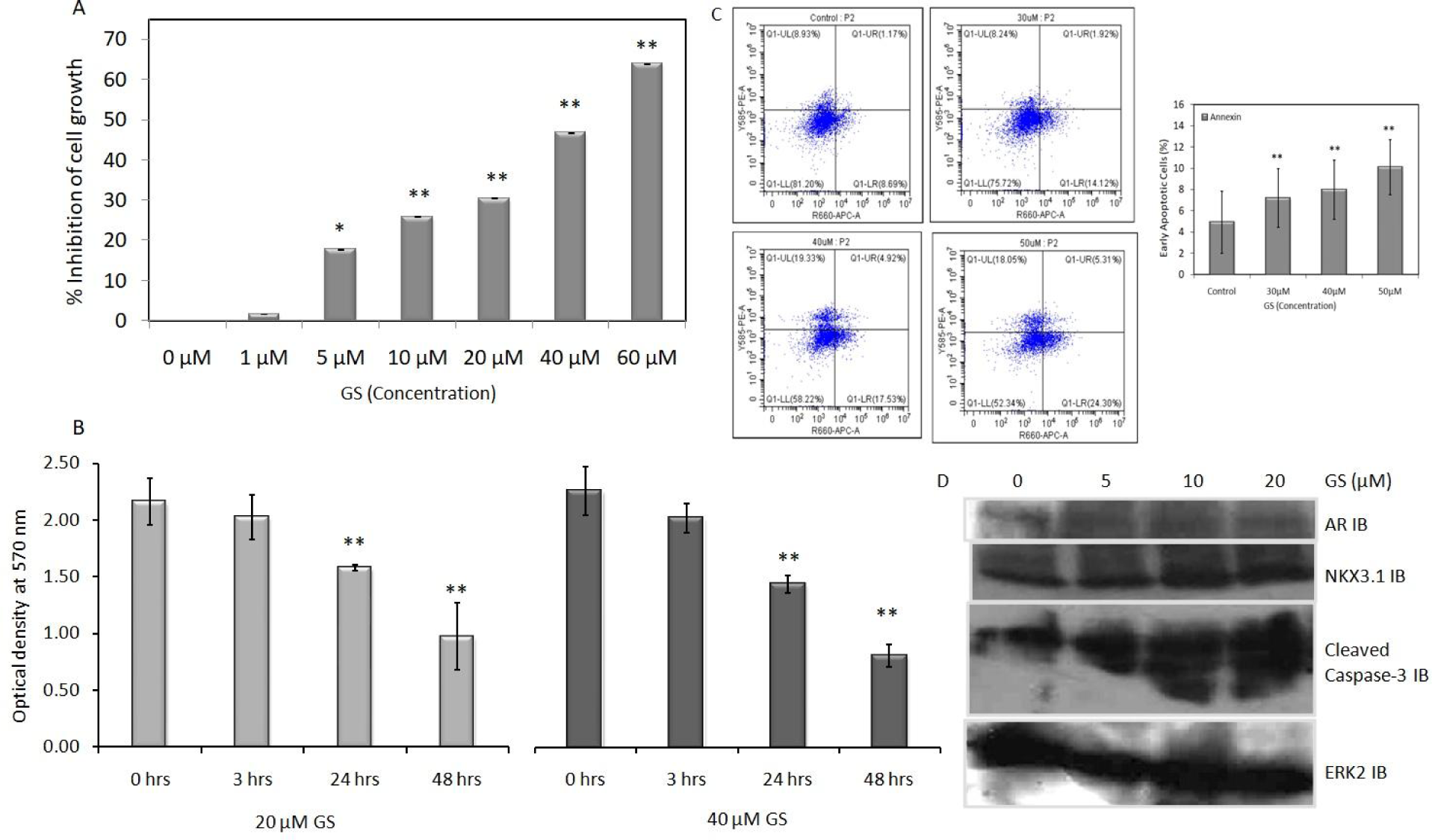
Effects of various doses of GS and time on viability and proliferation of LNCaP cells. (A) Human PCa LNCaP cells were seeded in a 96-well culture plate at a density of 3×105 cells/well and either treated with GS (1, 5, 10, 20, 40, and 60 μM) for 24 hours or vehicle control (0 μM). Inhibition of cell viability was measured using MTT assay. The optical density of vehicle control was taken as 0% inhibition and the corresponding fold change was plotted as % inhibition. (B) At ∼70% confluency, LNCaP cells were treated with GS (20 and 40 μM) for 3, 24, and 48 hours. Cell viability was measured using MTT assay. Optical density at 570nm was plotted as the mean±SD of the triplicate experiment. (C) The quantification of early/late apoptotic cells treated with the indicated concentration of GS (30, 40, and 50 µM) for 24 h by flow cytometry. The LNCaP cells were stained with Annexin V/FITC and propidium iodide. Q1UL: Necrosis, Q1UR: late apoptosis, Q1LL: viability, Q1LR: early apoptosis. At least 10,000 cells were analyzed per sample, and quadrant analysis was performed. Histogram showsmean±SD of early apoptotic cell percentage (n=3) *P<0.005, **P<0.001. (D) LNCaP cells were treated with GS for 24 hours, and whole-cell extracts were subjected to immunoblot analysis with the indicated antibodies. ERK2 was used as the loading control. Results are representative of three sets of independent experiments.

Temporal assessment of LNCaP cell viability was conducted at intervals of 0, 3, 24, and 48 hours, with GS concentrations of 20 and 40 µM. We observed a statistically significant decrement in cellular viability (p≤0.005) in LNCaP cells both at 24 and 48 hours at both 20 and 40 µM GS dosage (Figure 1B). Notably, PC3 cells only exhibited a statistically significant alteration in cellular viability after 48 hours of GS exposure (Supplementary Figures S1B and S1C).

Flow cytometry analyses, after GS treatment, disclosed a dose-proportional escalation in apoptotic cell fractions. Utilization of Annexin V staining facilitated the precise quantification of early-stage apoptotic cells. Post-24-hour GS exposure, a marked augmentation in early apoptotic cell counts was discernible in LNCaP cells subjected to 30, 40, and 50 µM GS concentrations, in comparison to vehicle-treated controls (Figure 1C). This affirms a dose-responsive induction of early apoptosis in LNCaP cells by GS.

Western blot assays were employed to evaluate the activation status of caspase-3, an enzyme pivotal to apoptotic pathways. Incremental elevations in cleaved caspase-3 levels were observed, corroborating the enzyme’s activation in a dose-dependent manner. Elevated Caspase-3 protein was observed even at the lowest examined GS concentration of 5 µM (Figure 1E).

In summary, our data substantiates the capacity of GS to initiate apoptosis in PCa cells in a dose and time-correlated manner. Cellular viability assays further corroborated these findings, particularly in LNCaP cells, as evinced by activated caspase-3 levels. These observations confirm GS as an anti-cancer agent, however warranting further exploration for the molecular basis of its pro-apoptotic capabilities.

### 3.2 Role of NKX3.1 in Modulating GS-Induced Apoptotic Responses

To elucidate the mechanistic role of NKX3.1 in GS-mediated apoptosis, we employed small interfering RNA (siRNA) to selectively inhibit NKX3.1 expression. Successful knockdown was confirmed through RT-PCR, illustrating a pronounced reduction in NKX3.1 transcript levels (Figure 2A).

**Figure 2:**
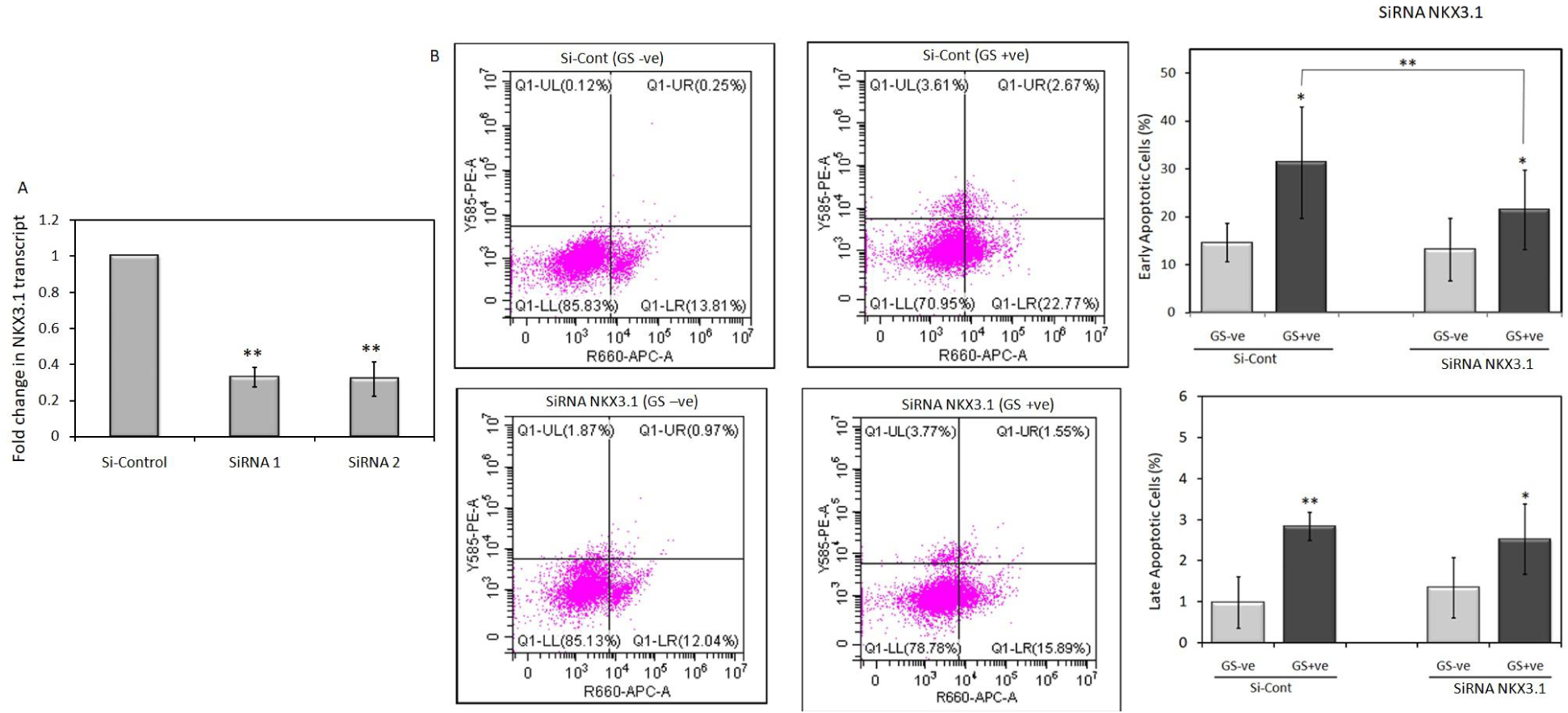
Effect of siRNA-mediated suppression of NKX3.1. (A) LNCaP cells were transfected with the indicated control siRNA or NKX3.1 -siRNA1 or -siRNA2. Four days after transfection cells were harvested, transcript levels of NKX3.1 were analyzed with real-time PCR. Histogram depicts fold change in transcript levels after siRNA transfection relative to Control siRNA and plotted as mean±SD. (B) LNCaP cells were transfected with control (Si-Cont) or NKX3.1 siRNA (siRNA NKX). 48 hours after transfection, cells were either left untreated (GS-ve) or treated with 40uM GS (GS+ve) for an additional 24 hours. For the quantification of early/late apoptotic cells by flow cytometry LNCaP cells were stained with Annexin V/FITC and propidium iodide. Q1UL: Necrosis, Q1UR: late apoptosis, Q1LL: viability, Q1LR: early apoptosis. At least 10,000 cells were analyzed per sample, and quadrant analysis was performed. Histogram depicts mean±SD of % cell numbers in the indicated quadrant with or without GS in siRNA control or siRNA NKX transfected cells, *P<0.005, **P<0.001.

Subsequently, we probed the influence of NKX3.1 knockdown on GS-mediated apoptotic induction. Flow cytometry analyses were conducted to quantify both early (Annexin V-positive) and late-stage apoptotic cells, juxtaposing these metrics in NKX3.1-inhibited and control cells following GS treatment. In the control siRNA (Si-Control) transfected cells treated with 40 µM GS, we documented a noteworthy escalation of 53% in early apoptotic cells; where the absence of GS produced 14.5% early apoptotic cells, GS exposure surged this number to 31.28%. Contrastingly, cells transfected with NKX3.1-targeted siRNA exhibited a more moderate increment, from 13.09% (GS –ve) to 21.37% (GS +ve), corresponding to a 38.7% rise. These observations intimate that NKX3.1 suppression significantly attenuates the apoptotic efficacy of GS (Figure 2B).

Further analyses revealed an analogous trend in late-stage apoptosis. In Si-Control transfected cells exposed to GS, the late apoptotic fraction elevated from 0.98% (GS –ve) to 2.84% (GS +ve), equating to a 65.57% increase. In sharp contrast, cells transfected with NKX3.1-specific siRNA manifested a less substantial increase, with late apoptotic cells rising from 1.34% (GS –ve) to 2.52% (GS +ve), marking a 46.96% increment. This consolidates the notion that NKX3.1 has a significant contributory role in facilitating GS-induced apoptosis, if not completely NKX3.1 dependent, and offers invaluable insights into the underlying molecular mechanisms. The implications of these findings are twofold: they not only advance our understanding of GS-induced apoptosis but also nominate NKX3.1 as a prospective therapeutic and/or prevention target in prostate cancer intervention strategies.

### 3.3 GS enhances NKX3.1 expression

To investigate the effects of GS on NKX3.1 expression in LNCaP cells, Western blot, and qRT-PCR analysis were performed following treatment with 20 and 40 µM of GS. The protein levels of NKX3.1 were evaluated at 24 hours and 48 hours of GS exposure. At 24 hours of GS exposure, the protein levels of NKX3.1 showed a significant increase, approximately twofold, compared to the control cells (Fig 3A-B). Notably, no significant changes in the protein levels of androgen receptor (AR) were observed at this time point, indicating that GS treatment specifically elevates NKX3.1 expression without affecting AR levels in LNCaP cells for up to 24 hours. Moreover, the effect of GS on NKX3.1 expression was sustained, showing persistent elevation for up to 48 hours, indicating a long-lasting response rather than an acute transient effect. However, with prolonged GS exposure for 48 hours, there was an induction of both NKX3.1 and AR protein expression in LNCaP cells, suggesting that extended GS exposure influences the expression of both genes. Considering this, subsequent studies were focused on the 24-hour time point (Figure 3A-C).

**Figure 3:**
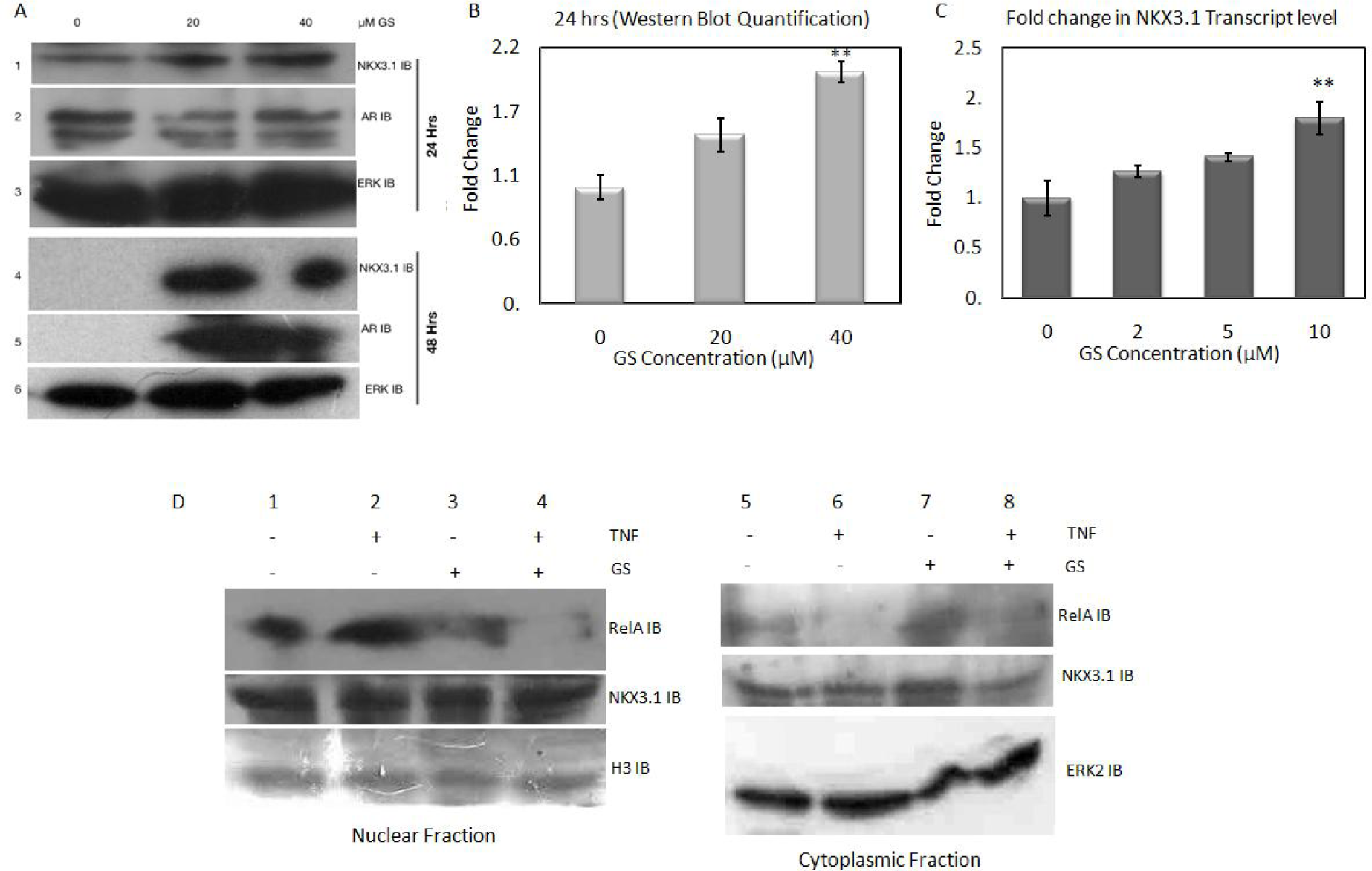
Effect of GS treatment on protein expression and localization: (A) LNCaP cells were treated with GS as indicated concentration. After indicated time periods, 24 or 48 hours, cells were harvested and whole cell extract was subjected to immunoblot analysis using specific antibodies. (B) Immunoblot bands were quantified using ImageJ. The histogram depicts fold change in NKX3.1 levels after 24 hours as compared with untreated samples. (C) The histogram depicts fold change in transcript levels measured using qRT-PCR after GS exposure at various indicated concentrations relative to vehicle-treated cells. (D) LNCaP cells were cultivated and either pre-exposed to GS (GS +) or left untreated (GS −) followed by TNFa for 20 minutes (TNF+) or not treated (TNF −). Subsequently, cells were harvested for nuclear and cytoplasmic proteins and were subjected to immunoblot analysis using antibodies. The nuclear translocation of RelA was determined with immunoblot analysis of nuclear fraction (left panel) and cytoplasmic fraction (right panel).

Since GS has established for its anti-inflammatory properties we analyzed both nuclear and cytoplasmic extract for the anti-inflammatory effects of GS, which is reflected by suppression of nuclear translocation of subunit ReL-A. GS induces NF-kB activity through IKK inhibition (25). LNCaP cells were pre-treated with or without 20 µM GS for 6 hours, followed by 20 minutes of TNFα treatment. Post-treatment cells were subjected to nuclear and cytoplasmic fraction isolation followed by immunoblot analysis of the RelA subunit in both fractions. TNFα induced NF-kB activation leading to RelA nuclear translocation (Fig, 3D, lane 2); we observed significantly low levels of RelA in nuclear fractions of cells exposed to GS and no localization into the nucleus after TNFα exposure which is supported by the consistent levels of RelA protein in the cytoplasmic fraction of GS-treated cells (Figure 3D, lane 7,8).

NKX3.1 protein and mRNA levels have been reported to be dramatically reduced by mitogens like EGF or PMA-ionomycin (P+I), TNFα, or IL-1α as early as one hour after stimulation (28). To explore the impact of GS on inflammation-induced NKX3.1 protein and transcript loss, LNCaP cells were exposed to mitogenic signals with either TNFα (Figure 4A) or PMA-Ionomycin (Figure 4B), in the presence or absence of GS. The results revealed that GS, despite being a known anti-inflammatory agent, was unable to rescue mitogen-induced NKX3.1 protein loss. The transcript level of NKX3.1 showed an expected elevation upon GS treatment (Fig. 4C), even when exposed to inflammatory signals. These findings suggest the potential involvement of different signaling axes in inflammation-induced NKX3.1 protein loss and GS-mediated NKX3.1 protein elevation (Figure 4).

**Figure 4:**
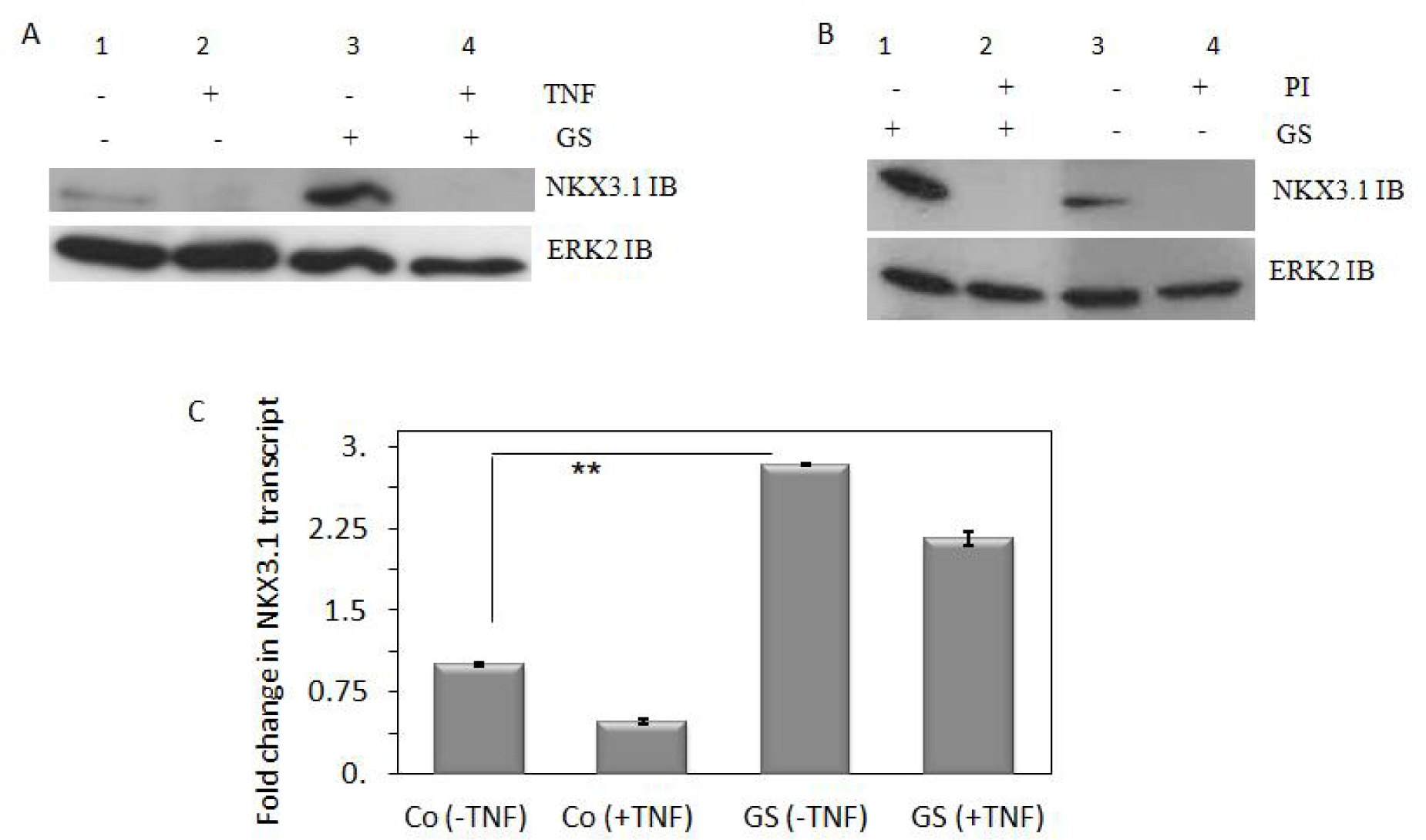
GS could not rescue TNFα (TNF) and PI (PMA+Ionomycine) induce NKX3.1 protein loss. Western blot analyses of whole cell extracts from LNCaP cells stimulated with either TNFα (A) or PI (B). Whole cell extracts were monitored with anti-NKX3.1 (upper panel), or internal control - anti-ERK2 (lower panel). (C) RTPCR analyses of NKX3.1 transcripts from LNCaP cells treated with TNFα. Bars depict fold change in NKX3.1 transcript levels when compared to Control, GS unexposed, and TNFα unstimulated, the presence (+) and absence (−) of each factor is depicted by + or −.

Since, NKX3.1 is one of the target genes of the Androgen receptor, we set out to understand the effect of hormone availability on GS-mediated effects on NKX3.1, levels of NKX3.1 protein and mRNA were measured in LNCaP cells cultured in charcoal-stripped serum (CSS) vs. hormone-supplied serum (NS) in the presence or absence of GS. Surprisingly, GS exposure not only increased NKX3.1 levels in standard media, as expected but also restored NKX3.1 protein in CSS, which had lost NKX3.1 due to the absence of hormones. The transcript levels of NKX3.1 also showed similar changes, reflecting the persistence of elevated NKX3.1 protein levels in these cells. qRT-PCR under the same condition shows a similar pattern of transcript expression. The absence of hormones led to the loss of NKX3.1 expression which was restored upon GS exposure (Figure 5A-B) both at protein and transcript level.

**Figure 5:**
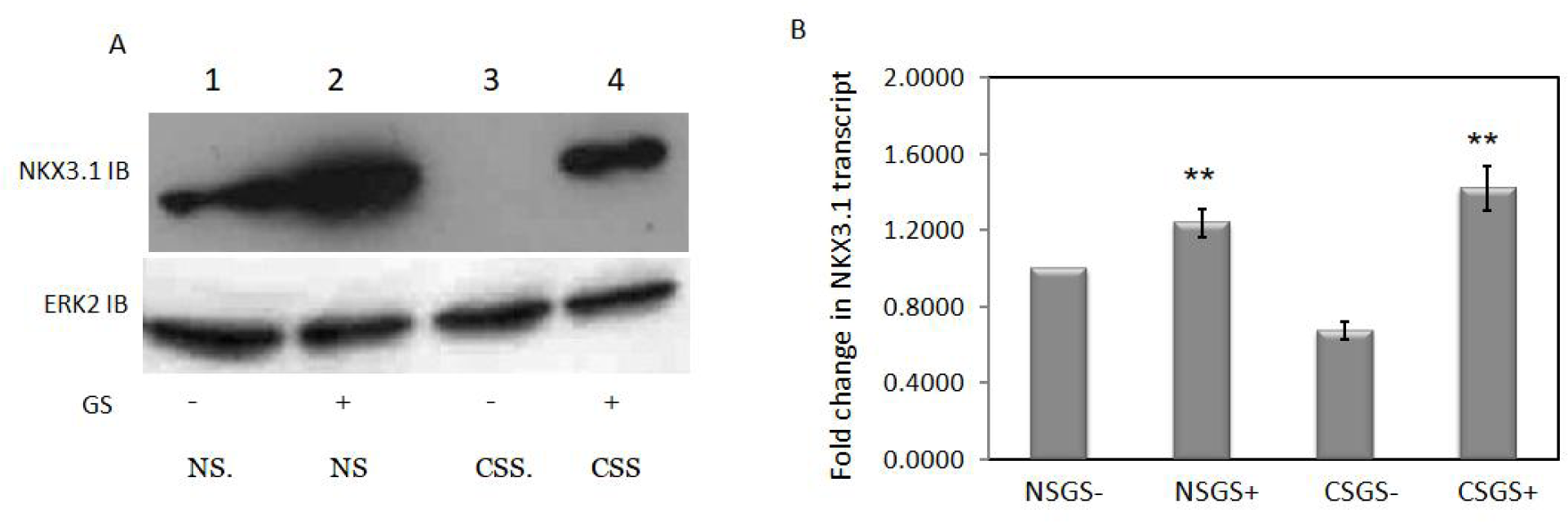
Effect of hormone-depleted media on GS-induced NKX3.1. LNCaP cells were cultivated in normally to reach 70% confluency in a 6-well plate. Later, media was either replaced with normal serum-supplemented RPMI (NS) or charcoal-stripped fetal calf serum-supplemented RPMI (CSS) for 2 hours before being exposed to GS (20 μM) for an additional 24 hours. Whole-cell protein extracts were subjected to immunoblot analysis using anti-NKX3.1 and anti-ERK2 antibodies. (A); along with RTPCR-based analysis for NKX3.1 Transcript levels. The histogram shows a fold change in transcript level as compared to control (NS without GS exposure) (B).

The results of this study indicate that GS enhances NKX3.1 expression in a time- and dose-dependent manner in LNCaP prostate cancer cells. GS-induced NKX3.1 expression was sustained over time, and GS exposure restored NKX3.1 protein levels even in the absence of hormones in the culture media, suggesting a potential role of GS in bypassing the AR dependence of hormone response genes. These findings highlight its potential as a modulator of NKX3.1 in prostate cancer cells.

To further elucidate the underlying mechanisms of GS-induced NKX3.1 expression, we aimed to identify its effect on the half-life (T_1/2_) of NKX3.1 at the protein (T_1/2_{protein}) and mRNA levels (T_1/2_{mRNA}). For the T_1/2_{mRNA}, LNCaP cells were treated with Actinomycin D (0.5 microgram/ml), a transcriptional inhibitor, and qRT-PCR analysis was conducted at various time intervals. LNCaP cells were treated with 40 μM of GS for 24 hours followed by Actinomycin-D treatment for different periods (0, 2, 4, 6, and 8 hours). ddCt-based fold change was considered for plotting a linear regression curve to calculate T_1/2_{mRNA} (Figure 6A). Without GS treatment, the T_1/2_{mRNA} was calculated to be 3.67 hours. Upon GS treatment, the T_1/2_{mRNA} was extended to approximately 4.53 hours, suggesting that GS enhances the stability of NKX3.1 mRNA by approximately 52 minutes (Figure 6A, histogram). This indicates that GS exerts its effect on the stability of NKX3.1 transcripts, resulting in a longer half-life.

**Figure 6:**
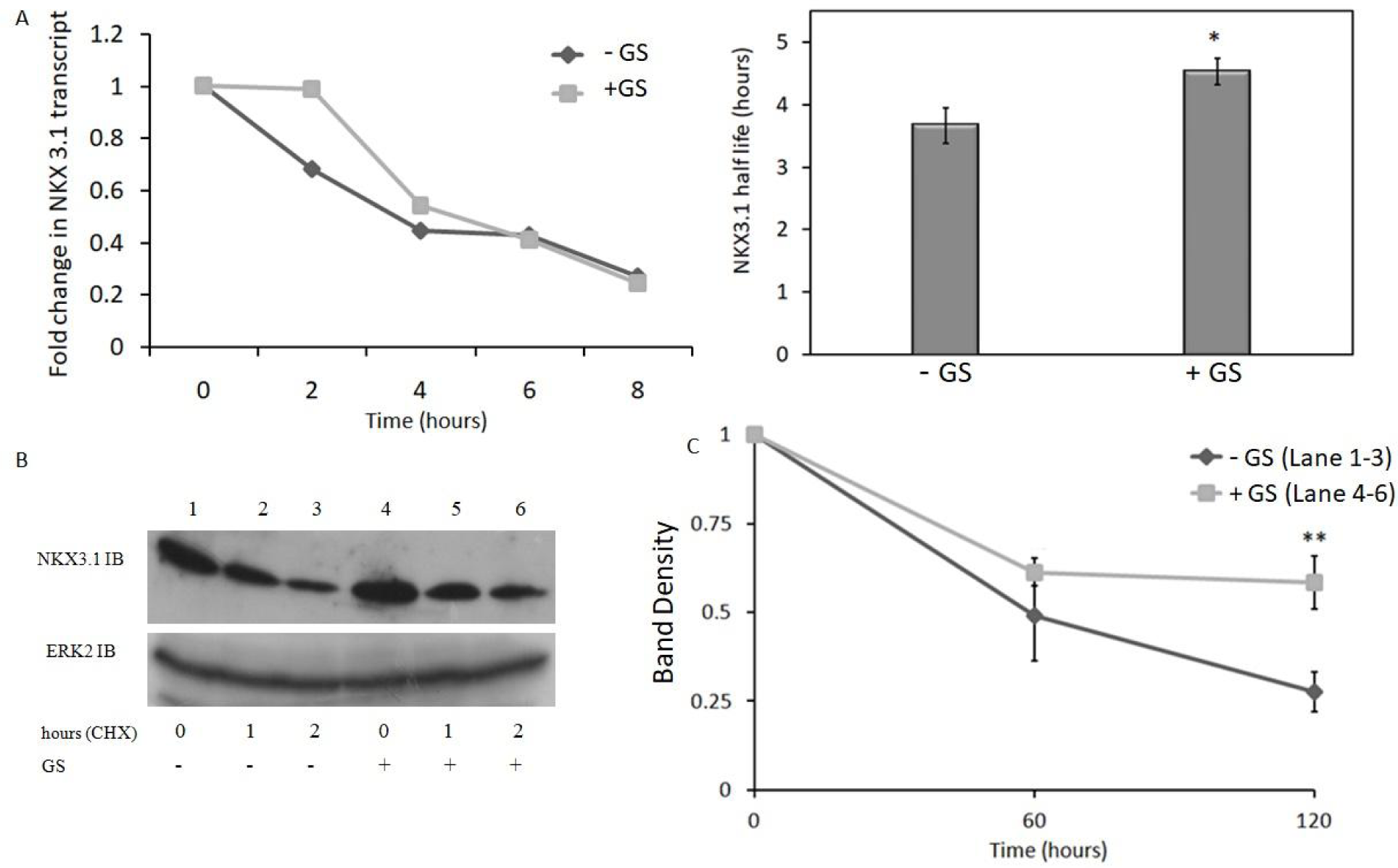
Effect of GS on NKX3.1 stability. Half-life (T_1/2_) of NKX3.1 protein (T_1/2_{protein}) and mRNA levels (T_1/2_{mRNA}) were calculated on a linear regression curve. (A) Cells were either preexposed with 40 μM GS or not exposed followed by actinomycin D treatment for 0, 2, 4, 6, and 8 hours. ddCt fold change data points were plotted as linear regression curve to calculate 0.5 fold change time point. Histogram depicts the mean±SD of T_1/2_{mRNA} calculated using 3 sets of experiments, *P<0.005, **P<0.001. (B) Cells were either preexposed with 40 μM GS or not exposed followed by Cycloheximide treatment for 0, 1, and 2 hours. After indicated hours cells were harvested for immunoblot analysis with indicated antibodies. Band intensities were normalized against control bands, and values were plotted on a linear regression curve.

For the protein stability assay (T_1/2_{protein}), LNCaP cells were treated with Cycloheximide to inhibit new protein synthesis, and Western blot analysis was performed at different time points. LNCaP cells were pre-treated with 40 µM GS (or vehicle as a control) for 6 hours before adding 10 µM cycloheximide to inhibit new protein synthesis. Sequential Western blot analysis of NKX3.1 levels was performed after 1 and 2 hours of cycloheximide treatment. Band density measurement was used to plot a linear regression curve to calculate T_1/2_{protein} with or without GS cultured cells (Figure 6B). The half-life of NKX3.1 protein without GS treatment was found to be approximately 74 minutes. In contrast, in the presence of GS, the T1/2 of NKX3.1 protein was approximately 127 minutes (Figure 6B, histogram). These results indicate that GS significantly affects the stability of NKX3.1 protein too.

No post-translational changes in NKX3.1 protein were visible in immunoblots following GS treatment. This suggests that GS primarily influences NKX3.1 expression by affecting mRNA and protein stability.

These findings provide valuable insights into the molecular mechanisms through which GS modulates NKX3.1 expression and highlight its potential as a preventive agent in the context of prostate cancer treatment. Further research is warranted to fully understand the signaling pathways involved in GS-induced changes in NKX3.1 stability and to explore the therapeutic implications of targeting NKX3.1 expression as a prostate cancer prevention strategy.

### 3.4 GS improves mitochondrial membrane potential

To assess the impact of GS treatment on mitochondrial membrane potential, we used Tetramethylrhodamine methyl esters (TMRM) staining, a fluorescent dye that specifically measures changes in mitochondrial membrane potential. The fluorescence intensity of TMRM is indicative of the health and functionality of mitochondria. LNCaP cells were treated with GS for 24 hours, and then the cells were collected to measure the intensity of TMRM fluorescence. The results of the TMRM staining demonstrated that GS treatment led to the increased fluorescence intensity of TMRM, indicating significantly enhanced mitochondrial membrane potential following 30 and 40 µM GS treatment (Figure 7).

**Figure 7:**
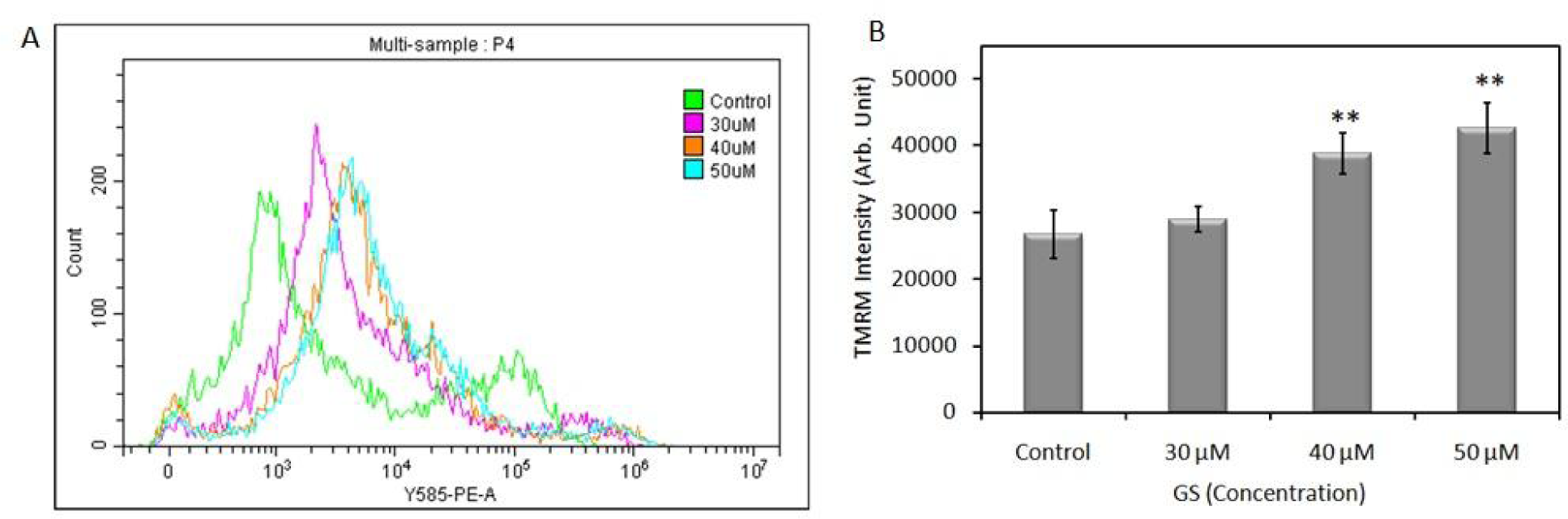
Mitochondrial membrane depolarization was measured using TMRM staining of LNCaP cells treated with GS (30,40, and 50 µM). Histogram Data are expressed as mean values ± sD (n=3) of TMRM intensity recorded for each event.

This suggests that GS treatment improved the mitochondrial membrane potential in LNCaP cells, indicating that GS may have beneficial effects on mitochondrial membrane integrity and cellular energy metabolism. Further research is warranted to fully understand the molecular mechanisms underlying the improvement of mitochondrial function by GS and to explore its potential implications for prostate cancer therapy and other mitochondrial-related disorders.

## Discussion

Prostate cancer is a highly heterogeneous and prevalent disease worldwide, with significant morbidity and mortality rates. Current treatment regimens for prostate cancer predominantly focus on androgen deprivation therapy (ADT), which often results in the emergence of castration-resistant prostate cancer (CRPC), leading to lethal outcomes. This makes it imperative to identify new targets beyond the androgen signaling pathway. In light of these challenges, our study showcases the potential of GS as a preventive agent in PCa by elucidating its anticancer mechanism. The focus on NKX3.1 explores an alternative pathway that may act independently or synergistically with androgen deprivation therapy.

Guggulsterone (GS) has garnered attention for its anti-carcinogenic properties, with numerous studies demonstrating its antiproliferative, anti-metastatic, anti-angiogenic, and pro-apoptotic effects in various cancer cell lines (26) and animal models. However, the underlying mechanisms and key events leading to the anti-cancer effects of GS still need to be completely understood. GS is known to modulate the activity of nuclear receptors, growth factors, and transcription factors, specifically, z-GS was reported to be the antagonist of the androgen receptor (27). Our findings present compelling evidence that guggulsterone (GS) upregulates NKX3.1 expression at both the protein and mRNA levels, a particularly significant observation given that NKX3.1 is a known androgen receptor target. Intriguingly, this upregulation was maintained even under hormonal depletion, thereby suggesting a mechanism that circumvents the conventional androgen receptor (AR) dependency for NKX3.1.

The sustained expression of NKX3.1 in hormone-depleted conditions posits GS as a potential agent that can act either in conjunction with or independently of androgen deprivation therapy (ADT). GS effects the stability of NKX3.1 mRNA and protein, contributing to the sustained increase in NKX3.1 expression. The increased NKX3.1 levels might contribute to increased mitochondrial stability, leading to protective effects against oxidative stress. Oxidative stress is one of the critical factors in cancer initiation and progression brings forth intriguing implications that tie in with the recent discovery of NKX3.1’s role in mitochondrial function (29). NKX3.1 has been reported to serve as a modulator of cellular energy homeostasis through oxidative phosphorylation by regulating the transcription of mitochondrial-encoded ETC genes (30). In concordance between the stabilization of mitochondrial membrane potential and upregulated NKX3.1 expression in GS-treated cells could potentially point towards an integrative mechanism where GS’s anticancer activities are mediated, at least in part, through the fortification of mitochondrial function via NKX3.1. Moreover, our data indicates a functional link between GS-induced apoptosis and NKX3.1 levels in prostate cancer cells. When NKX3.1 expression was knocked down, the pro-apoptotic effects of GS were substantially mitigated, corroborating that NKX3.1 is an integral part of the GS-mediated apoptotic pathway. The observation that elevated NKX3.1 levels, induced by GS treatment, are positively correlated with increased mitochondrial membrane potential. It opens up avenues for future research to explore the connections between GS, NKX3.1, and mitochondrial function in more nuanced models.

Thus, the NKX3.1-GS correlation is not merely an attractive target for preventive intervention, but also a prospective entry point for understanding the interconnectedness of cellular energy metabolism, oxidative stress, and prostate cancer pathophysiology.

Overall, our research adds to the growing understanding of the diverse molecular mechanisms through which GS exerts its anti-cancer effects on prostate cancer cells. Further investigations into the signaling pathways and mechanisms underlying GS-induced changes in NKX3.1 expression and apoptosis are warranted.

The results of this study open up new avenues of research for the development of targeted therapies that stabilize NKX3.1 expression and exploit its role in protecting mitochondria, thus offering promising prospects for prostate cancer chemoprevention and treatment strategies.

## Conclusion

In conclusion, We demonstrated that GS induces apoptosis in prostate cancer cells, with a dose-dependent and time dependent manner. Our research highlights the critical role of NKX3.1 in mediating GS-induced apoptosis. Our study revealed that elevated NKX3.1 levels in GS-treated cells are associated with increased mitochondrial membrane potential, potentially suppressing prostate cancer by protecting mitochondrial function. Conclusively, our study adds to the growing body of evidence supporting the potential of GS as a chemopreventive and therapeutic agent for prostate cancer. It encourages further exploration of the importance of NKX3.1 as a potential therapeutic or preventive target in prostate cancer. We hope that our findings will inspire further investigations and pave the way for future clinical trials to assess the efficacy of GS-based treatments in prostate cancer patients, bringing us closer to more effective and tailored therapies.

## Funding

This work was supported by the Department of Science and Technology Women Scientist-A scheme, Institute of Eminence, BHU (MPDF to GJ), Banaras Hindu University; (RET Non-Net Fellowship to N., PR, CD).

## Conflict of Interest Statement

The authors have no conflicts of interest to declare.

---

**Supplementary Figure 1:**
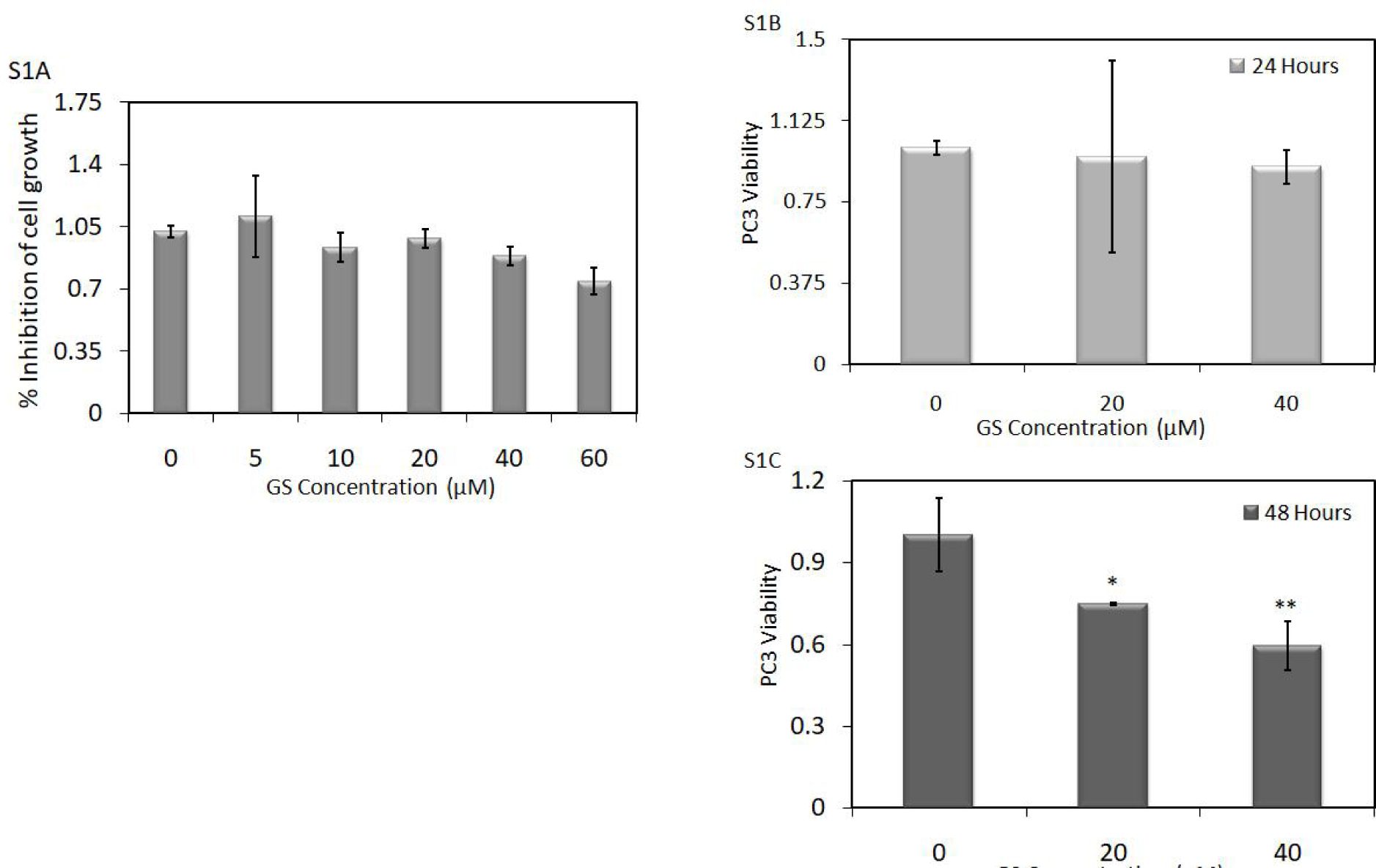
S1: Effects of various doses of GS and time on viability and proliferation of PC3 cells. (A) Human PCa PC3 cells were seeded in a 96-well culture plate at a density of 3×105 cells/well and either treated with GS (5, 10, 20, 40, and 60 μM) for 24 hours or vehicle control (0 μM). Inhibition of cell viability was measured using MTT assay. The optical density of vehicle control was taken as 100% growth and the corresponding fold change was plotted. At ∼70% confluency, PC3 cells were treated with GS (20 and 40 μM) for 24 (B), and 48 hours (C). Cell viability was measured using MTT assay. Cell viability was plotted as the mean±SD of the triplicate experiment.

## Notes

### Competing Interest Statement

The authors have declared no competing interest.

### Summary of Updates

The sequence of authors is mismatched in PDF and Author Information section.

