## Supplementary figures and images for "Guggulsterone Enhances NKX3.1 Expression and Induces Apoptosis in Prostate Cancer Cells: Implications for Chemoprevention"

### Supplementary Figure 1

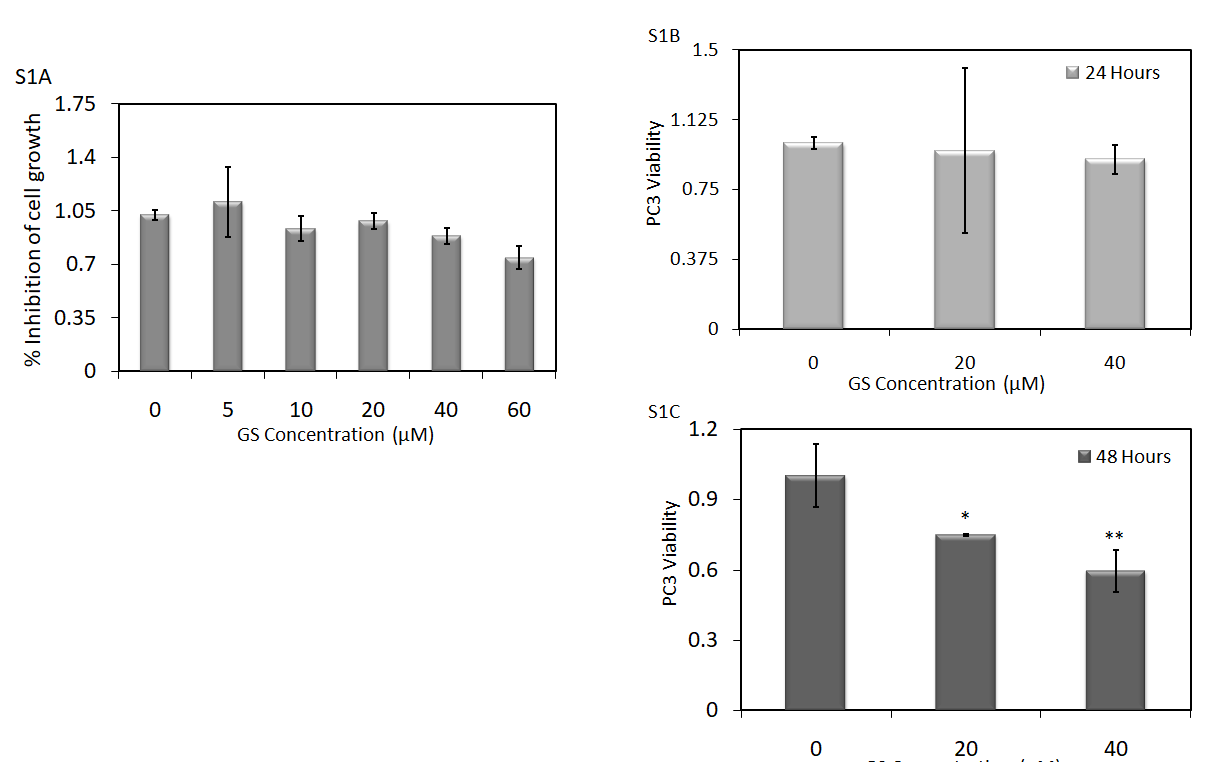
